# Genome-wide association study of HIV whole genome sequences validated using drug resistance

**DOI:** 10.1101/076216

**Authors:** Robert A. Power, Siva Davaniah, Anne Derache, Eduan Wilkinson, Frank Tanser, Ravindra K. Gupta, Deenan Pillay, Tulio de Oliveira

## Abstract

Genome-wide association studies (GWAS) have considerably advanced our understanding of human traits and diseases. With the increasing availability of whole genome sequences (WGS) for pathogens, it is important to establish whether GWAS of viral genomes could reveal important biological insights. Here we perform the first proof of concept viral GWAS examining drug resistance (DR), a phenotype with well understood genetics.

We performed a GWAS of DR in a sample of 343 HIV subtype C patients failing 1^st^ line antiretroviral treatment in rural KwaZulu-Natal, South Africa. The majority and minority variants within each sequence were called using PILON, and GWAS was performed within PLINK. HIV WGS from patients failing on different antiretroviral treatments were compared to sequences derived from individuals naive to the respective treatment.

GWAS methodology was validated by identifying five associations on a genetic level that led to amino acid changes known to cause DR. Further, we highlighted the ability of GWAS to identify epistatic effects, identifying two replicable variants within amino acid 68 of the reverse transcriptase protein previously described as potential fitness compensatory mutations. A possible additional DR variant within amino acid 91 of the matrix region of the Gag protein was associated with tenofovir failure, highlighting the ability of GWAS to identify variants outside classical candidate genes. Our results also suggest a polygenic component to DR.

These results validate the applicability of GWAS to HIV WGS data even in relative small samples, and emphasise how high throughput sequencing can provide novel and clinically relevant insights. Further they suggested that for viruses like HIV, population structure was only minor concern compared to that seen in bacteria or parasite GWAS. Given the small genome length and reduced burden for multiple testing, this makes HIV an ideal candidate for GWAS.

## Introduction

Genome-wide association studies (GWAS) have led to significant advances in the understanding of complex human traits and diseases. They involve the analysis of hundreds of thousands or millions of common genetic variants, usually single nucleotide polymorphisms (SNPs), testing for an association between each variant and a phenotype (see [1]). This allows for the analysis of many variants across the genome, blind to their location or functionality. This approach has identified hundreds of causal risk variants for dozens of diseases in the last decade (e.g. [2–4]), each a potential drug target for novel treatments. These advances were made possible due to the availability of cost effective SNP genotyping technology which capture known common genetic variants. The limitation of this approach is that it misses variants absent from the chip, especially rare or *de novo*mutations. For this reason, genetic research is increasingly moving towards whole genome sequencing approaches to capture the full range of genetic variants in a population.

In this respect, the field of pathogen genomics is quickly catching up with human genomics, with international collaborations currently generating thousands of whole genome sequences (WGS) for pathogens such as HIV and malaria (e.g. the PANGEA Consortium[5] and the MalariaGen Consortium[6]). These WGS allow for the application of GWAS-style identification of novel genetic risk variants without the need for SNP genotyping chips.

A GWAS approach has previously been successfully applied to other non-virus pathogen, almost always using treatment resistance or failure as the phenotype[7]. These studies have included *Plasmodium falciparum*[8], *Mycobacterium tuberculosis*[9], *Staphylococcus aureus*[10] and *Streptococcus pneumoniae*[11]. Sample sizes have ranged from 75 to 3,701 sequences, and in even smaller samples have identified both novel and known variants that capture almost all the variation in treatment outcome.

However it is still unclear how well suited the viral genome is to a GWAS approach. The only viral GWAS to date combined GWAS of human SNP and HIV amino acid data, and identified multiple host genetic variants in the HLA region associated with HIV amino acid diversity[12]. However they found no associations between the HIV genome and their outcome of interest, viral load. The high percentage of coding sequence in viral genomes and overlapping reading frames may constrain the polygenic architecture for which GWAS was conceived: with many variants each of individually small effect. Another limitation of previous studies was that they did not allow for heterozygosity. Heterozygosity at a locus can arise due to mixed infections or within-host pathogen genetic diversification. Although this is rare in most pathogens studied with GWAS to date, it is highly relevant to many viral infections. Lastly, parasite and bacterial GWAS have observed a large level of population structure presumably due in part to homologous recombination and recent selection[13]. Given the challenges faced by previous analyses, more work is needed to properly define the genomic architecture of viruses and whether it is suitable to a GWAS style approach.

To validate the effectiveness of a viral GWAS we aimed to replicate the success of bacterial GWAS and focus on a phenotype known to be under strong selection pressure, specifically antiretroviral therapy (ART) resistance in HIV. The provision of ART to over 6.2 million people living with HIV in sub-Saharan Africa has been one of the most successful public health interventions ever undertaken[14], improving life expectancy[15], and reducing transmission[16, 17]. As a result, ART has been one of the most potent selection pressures on HIV. Given its importance to global health, resistance to ART has been well studied in HIV with many amino acid changes known to lead to DR [18]. Thus, DR is a useful phenotype for validating GWAS in HIV as findings can be compared to the existing literature as well as to large publically available databases of genes involved in HIV DR. In this study, we aim to identify known variants and validate the applicability of GWAS methods to the HIV genome.

## Results

### Genomic architecture of HIV SNPs

343 samples with phenotype and genotype data remained after variant calling and quality control (Table 1). A total of the 5379 SNPs with a minor allele frequency>= 1% were identified. An excess of rare variants was observed with a mean allele frequency of 11.3% and median of 6.0% (see Supplementary Figure 1). Additionally 2502 variants were identified with a frequency less than 1% though not included in the analyses. Variants were evenly distributed across the genome, despite missingness differing by region (see Supplementary Figure 2). The permuted threshold for genome-wide significance was p=7E-5, less stringent than that derived by Bonferroni correction for the number of variants (p=9.3E-6) and suggesting that there was substantial correlation between SNPs. This correlation is expected, due to the close proximity of SNPs in WGS data which leads to linkage disequilibrium and the non-independence of tests. As such, genome-wide significance was determined using the permutation adjusted p-permutation adjusted p-value threshold. SNPs were labelled by their base pair position plus reference allele, e.g. 1A. SNPs were also linked to their corresponding amino acid position in the different HIV proteins using reference sequence AF411967.

**Table 1:** Number of WGS treated with each drug, and correlations between drugs within samples

| Drug | Treated | Untreated | Correlation with: |  |  |  |  |  |
| --- | --- | --- | --- | --- | --- | --- | --- | --- |
|  |  |  | Zidovudine | Stavudine | Tenofovir | Efavirenz | Nevirapine | Lopinavir |
| **Zidovudine** | 32 | 311 | 1 | - | - | - | - | - |
| **Stavudine** | 291 | 52 | -0.058 | 1 | - | - | - | - |
| **Tenofovir** | 101 | 242 | -0.117 | -0.507 | 1 | - | - | - |
| **Efavirenz** | 259 | 84 | 0.011 | -0.023 | 0.128 | 1 | - | - |
| **Nevirapine** | 113 | 230 | -0.017 | 0.127 | -0.057 | -0.623 | 1 | - |
| **Lopinavir** | 26 | 317 | 0.213 | -0.033 | 0.053 | -0.151 | -0.115 | 1 |

**Figure 1:**
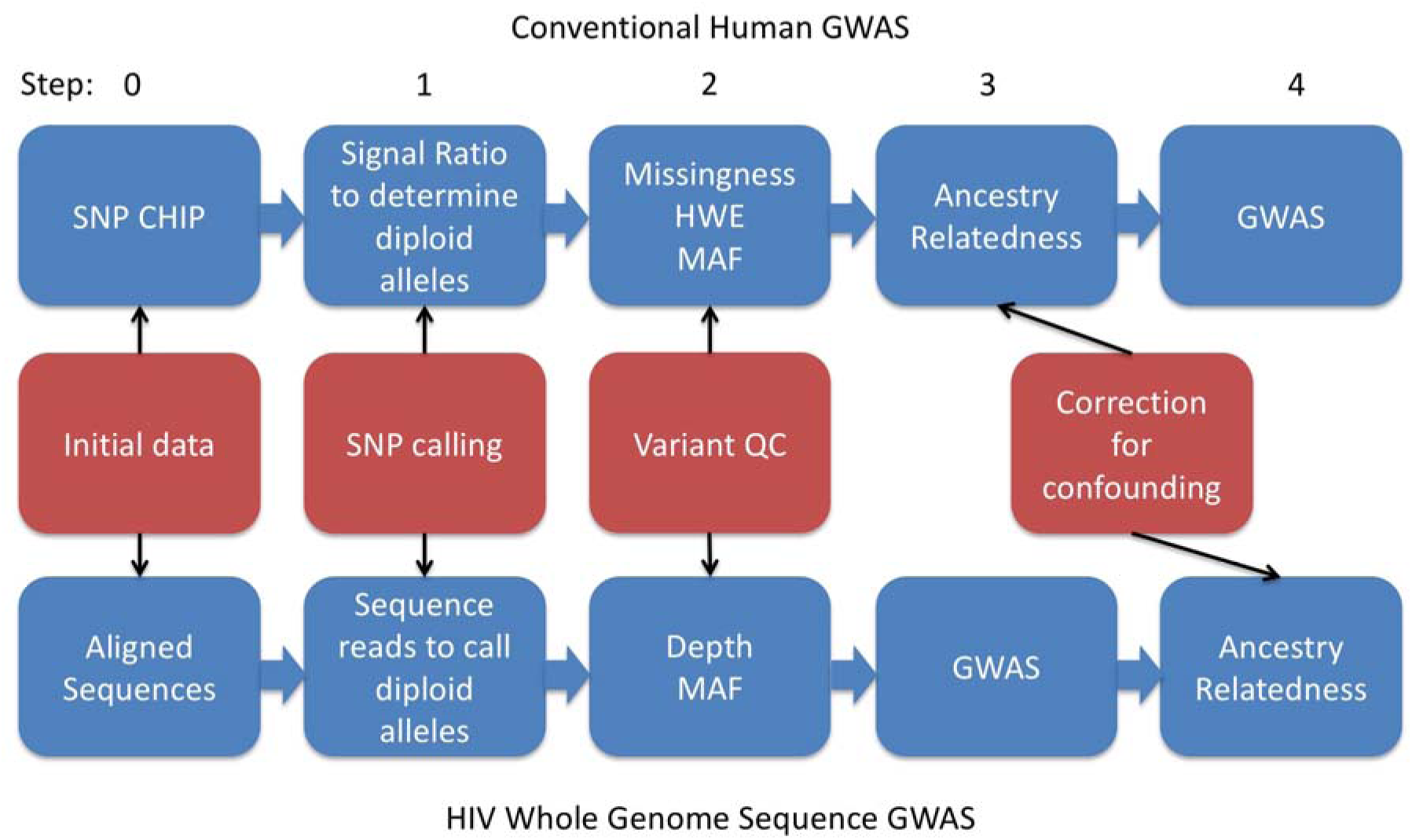
Analysis pipeline for HIV whole genome sequence (WGS) genome-wide association study (GWAS) compared to a human study using a SNP chip. Step 1) Diploidy defined for both human and pathogen, to reflect ‘real’ heterozygosity and heterozygosity from within host viral diversity. 2) While missingness and Hardy-Weinberg Equilibrium are used to assess genotyping quality in human GWAS, in viral GWAS we used depth of sequencing to assess variant calls. As such, higher calling confidence is associated with higher missingness in viral SNPs, while the reverse is true in humans. Low minor allele frequency (MAF) is always used to remove variants that have low power to detect effects and may reflect errors. 3&4) Correction for ancestry and relatedness are key to human GWAS, however due to both more homogenous sampling and difficulty in applying conventional corrections in human data to viral, this was done as a sensitivity test in a smaller sample for top SNPs in HIV GWAS.

**Figure 2:**
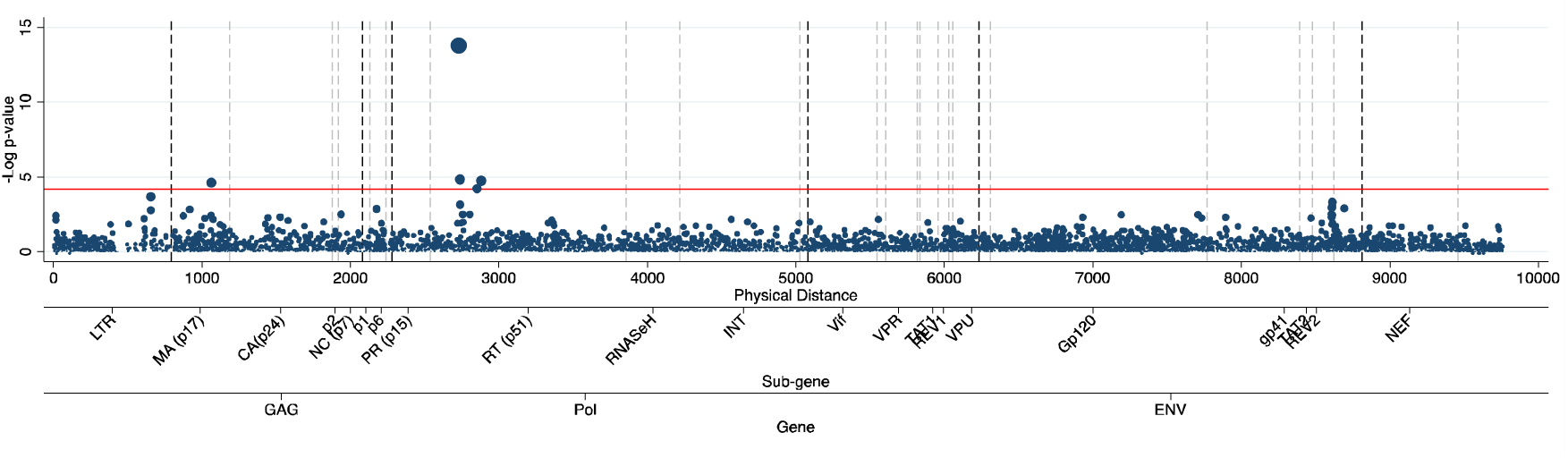
Manhattan plot comparing HIV sequences that were exposed to tenofovir to those that were not. The reference line at p=7E-5 is the line for permutation adjusted genome wide significance. Dashed grey lines on genomic locations refer to borders of genes (black dashed refer to GAG, Pol and ENV). Each marker is a SNP, weighted by it’s –log(p-value) to highlight the most significant SNPs.

### Validating GWAS with known DR variants

GWAS was performed to identify variants associated with drug resistance. The drug resistance phenotype was binary for each drug and defined as any history, or not, of failure while treated with the given drug. Failure was defined as at least one measure of viral load >1000 copies/ml after 12 months of treatment. GWAS identified eight independent associations at permutation adjusted genome-wide significance. Five of the associations were known loci involved in DR and all but one were in the reverse transcriptase region (RT), the functional target of these drugs (see Table 2). Failure on tenofovir was associated with three known SNPs (2730G, 2852A and 2880T, see Figure 1) in the RT region, at amino acid positions 65, 106, and 115, of which position 65 and 115 were known tenofovir DR variants and position 106 was previously associated with DR with the most common drugs used in combination with tenofovir. Treatment with zidovudine was associated with SNP 2745G, a known drug variant in RT amino acid 70 (Supplementary Figure 3). Nevirapine treatment was associated with a SNP (3078G) at RT 181, again a previously known DR variant (Supplementary Figure 4). No associations were seen with known resistance variants for lopinavir, efavirenz and stavudine (Supplementary Figure 5, 6 and 7). These results remained significant after correction for confounding from population structure and length of treatment (Supplementary Table 1)

**Table 2:** Results for genome-wide significant SNPs and their corresponding amino acid positions. Note that the effect of SNP 2739A is protective against stavudine resistance (i.e. odds ratio [OR] <1) and the association is actually with tenofovir, that has a negatively correlated prescription regime. Ref.=Reference; BP= base position; A1 = effect allele; Cis=proximal to known DR variant; Conv.=convergent, i.e. known R variant for another drug; OR=Odds ratio; SE=standard error.

| Drug | SNP | Missing<br>-ness | A1 | Ref. | Gene | Amino<br>acid N | Ref. Amino<br>Acid | A1 Amino<br>Acid | Known | OR | SE | Unadjust<br>ed p-value | Permutatio<br>n adjusted<br>p-value |
| --- | --- | --- | --- | --- | --- | --- | --- | --- | --- | --- | --- | --- | --- |
| Nevirapine | 3078G | 14% | G | A | RT | 181 | Y | C | Yes | 5.20 | 0.26 | 4.77E-10 | 1.00E-07 |
| Stavudine | 2739A | 14% | A | G | RT | 68 | S | N | Cis | 0.08 | 0.54 | 5.38E-06 | 0.0081 |
| Tenofovir | 1063A | 18% | G | A | MA (p17) | 91 | R | G | No | 1.79 | 0.14 | 2.42E-05 | 0.016 |
|  | 2730G | 13% | G | A | RT | 65 | K | R | Yes | 6.44 | 0.24 | 1.67E-14 | 1.00E-07 |
|  | 2738G | 14% | G | A | RT | 68 | S | G | Cis | 2.89 | 0.24 | 1.45E-05 | 0.0088 |
|  | 2852A | 14% | A | G | RT | 106 | V | M | Conv. | 1.72 | 0.14 | 6.19E-05 | 0.047 |
|  | 2880T | 13% | T | A | RT | 115 | Y | F | Conv. | 5.77 | 0.41 | 1.80E-05 | 0.011 |
| Zidovudine | 2745G | 16% | G | A | RT | 70 | K | R | Yes | 3.11 | 0.22 | 2.94E-07 | 0.0006 |

**Figure 3:**
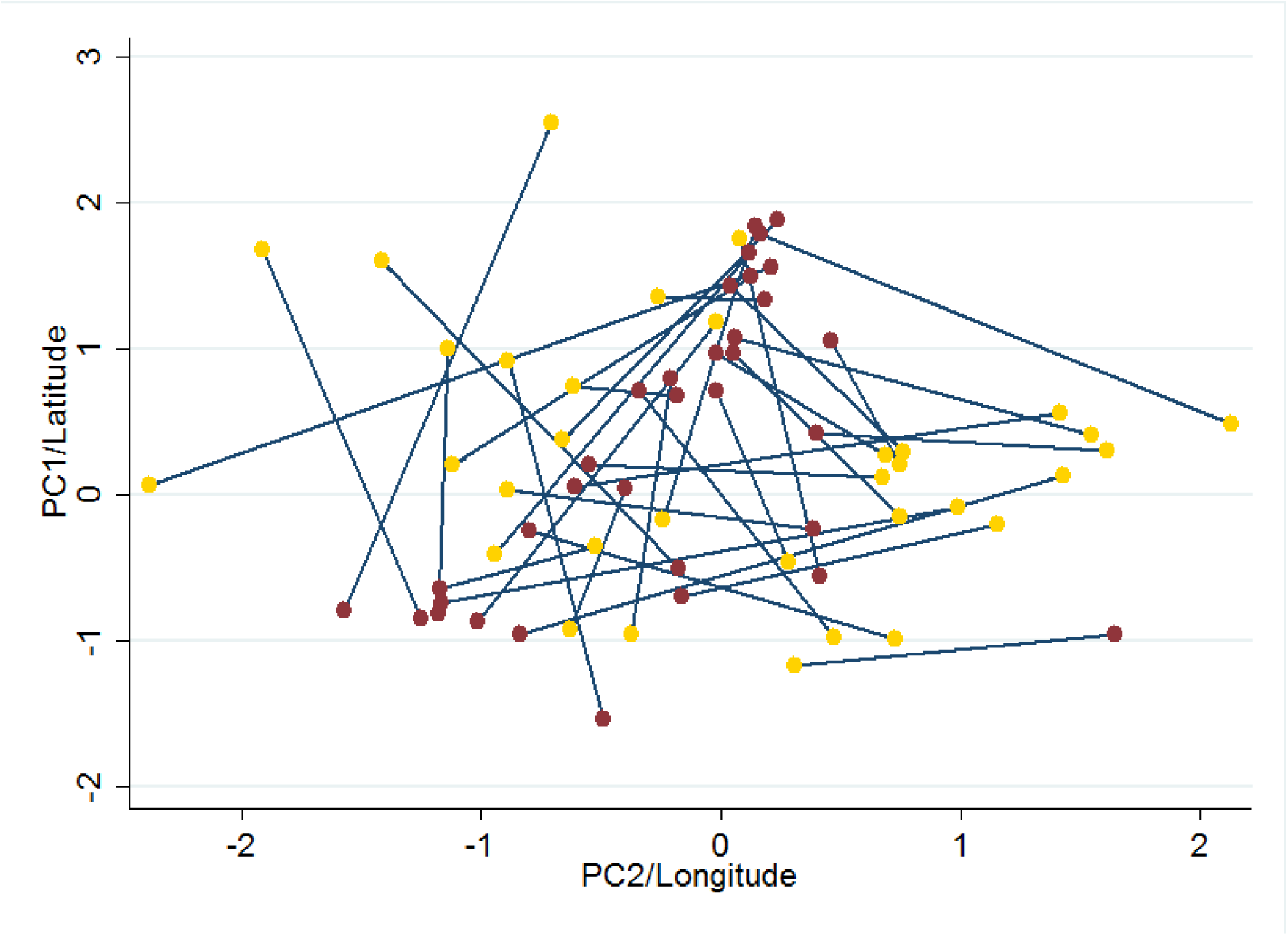
Plot of standardised values for the ancestry informative principle components 1 & 2 (red) and latitude & longitude (gold) for HIV sequences, with values for each sequence linked by a line. No correlation between geographic position and genetic position was observed.

While our analyses identified several known variants for DR, not all were identified. However, it is well known in GWAS studies that sample size is a critical limitation, with additional SNPs identified when larger samples are available. We observed a weak positive correlation in our analyses between the number of significant associations per drug and sample size (R^2^=22%). Looking at known DR mutations[18] with data available showed an excess of significant associations compared to expectation by chance, with 12% containing a variant at genome-wide significance and a further 41% containing at least one at nominal significance, despite incomplete coverage (p<0.001; see Supplementary Table 2). This trend was especially clear within the primary resistance mutations.

### Identification of novel variants

As well as known drug resistance variants, additional associations were observed. The first was two associations within RT amino acid 68. The first was between tenofovir failure and SNP 2738G resulting in a change from serine to glycine. Replication was performed using the Stanford University HIV Drug Resistance Database[19, 20]. Subtype C sequences within the Stanford database from individuals failing to tenofovir or other NRTIs (n=9,357) all had the reference (serine) amino acid. For sequences showing resistance to tenofovir, however, 5.5% had glycine at this position (n=488, p<0.0001 compared to non exposed distribution). Stavudine resistance was showed associations with a different SNP (2739A) from serine to asparagine (Supplementary Figure 3). However further investigation showed this to be an association with the negatively correlated drug tenofovir which had a p-value just below genome-wide significance for this SNP (p=7.1E-4). This was clear both from the fact the reference sequence allele was associated with stavudine DR, and from the results of the replication. For sequences failing on tenofovir 4.7% had asparagine at position 68 (p<0.0001), while for sequences failing on stavudine (n=2,800) no asparagine variants were observed. While not known drug resistance variants, amino acid 68 (specifically the change to glycine) has been suggested as a compensatory mutation for reduced fitness due to the drug resistance variant in amino acid 65[21, 22]. Indeed epistasis was observed between the significant SNPs in amino acid 68 and those in amino acids 65 and 106 (Supplementary Table 3).

For tenofovir failure an association was also seen with SNP 1063A in amino acid 91 of the matrix region, an entirely novel association (see Figure 2). Whilst not available in the Stanford database, we compared our results to the Los Alamos HIV Sequence Database drug naïve WGS at the amino acid 91 of the matrix region. Interestingly, for the amino acid 91 we observed a high level of genetic variation, with coding for nine amino acids. Focusing on the associated genetic change, we observed significantly different (p<0.0001) frequencies in the drug naïve sample (37% G vs. 61% A) compared to the tenofovir-exposed sequences in our sample (65% G vs. 35% A). Our WGS tenofovir naïve cases had a same frequency as the publically available sequences (37% G). While not an independent replication, this lends some support to our finding.

### Population stratification and cryptic relatedness

A concern in GWAS is the possibility of confounding by population stratification, which can lead to a systematic inflation in the number of false positives. QQ plots are a standard tool for testing for inflation in GWAS, plotting observed p-values across the genome compared to expected p-value distribution. These suggested a systematic deflation in p-values in this study, with genomic lambdas between 0.66-0.80. The lambda value is derived from the median observed chi squared statistic divided by the median expected chi squared statistic (for p=0.5). Under the null distribution, a lambda of 1 is expected, with a value above 1.05–1.10 usually taken as evidence of inflation. However, the excess of very rare variants (see Supplementary Figure 1) prevented a normal distribution of p-values, with a reduced number of significant SNPs compared to expected under the null. Restricting the analysis to SNPs where minor allele frequency was at least 10% supported this hypothesis, with an increase in genomic lambdas (0.81–1.00). To account for this, we compared our distribution of p-values to those when the phenotype was permuted within our data. This removed the systematic deflation in our expected vs. observed p-value distributions (lambdas 0.99–1.36, median 1.076), now showing a distribution close to null for the majority of SNPs (see Supplementary Figures 8–13). An inflation of p-values compared to permuted phenotypes was observed only within the tail end of highly significant SNPs. This is a characteristic not of population stratification but of a trait being polygenic, i.e. with many truly causal SNPs each explaining only a small proportion of variance. This distribution is common among human GWAS QQ plots and suggests larger studies of DR will yield additional causal SNPs, albeit with smaller effect sizes.

Usually population stratification is addressed by correcting for ancestry informative principal components. These principal components are based on SNP correlations across the genome, and have been shown to accurately capture population structure[23]. However their construction proved difficult in our total sample due to much higher missingness than is typical in GWAS data from genotyping chips. As such we performed a sensitivity analysis in a smaller sample with near complete sequencing (n=178) to test the effect of our genome-wide significant SNPs after correcting for principal components. No large attenuation of effect was observed, with half of the genome-wide significant SNPs showing an increased effect size when the first five principal components were included as covariates (Supplementary Table 1). Predictably we observed higher p-values in the sensitivity test due to the much smaller sample size. The partial availability of GPS data for individual’s household allowed for comparison of geographic proximity to genetic similarity (n=34). We did not observe clear genetic clustering overlapping with geographic (see Figure 3), though a pairwise comparison of genetic distance based on coordinates of first 2 principal components and geographic position did show a weak association between the two (R^2^=1.4%, p<0.005).

Another potential confounder within GWAS is relatedness between samples. Traditional measures of human relatedness were not appropriate for the analysis of pathogen genomics data. We performed a sensitivity test to remove samples closely linked within phylogenetic clusters (N=6). The results did not differ greatly, suggesting our top findings were not driven by population stratification or cryptic relatedness (see Supplementary Table 1).

## Discussion

In this study, we performed a proof of concept analysis that shows how a GWAS approach can identify many known variants and replicable novel associations using HIV WGS. We identified five variants at loci which corresponded with amino acid changes previously associated with DR. While not all previously known DR variants were identified at genome-wide significance in our analyses, we observed an excess of nominally significant associations at these loci (p<0.001, Supplementary Table 2). This is reminiscent of the polygenicity observed in human GWAS. Often an excess of sub-genome-wide significant variants was identified prior to identifying those specific SNPs truly associated with a trait[24]. We can expect many of those previously known variants to become genome-wide significant once sample sizes increase.

As well as validating known variants, our results highlight two ways in which GWAS can identify potential novel variants. The first is by identifying variants of smaller or indirect effects, such as via epistasis. We identified two nonsynonymous variants changing the RT amino acid 68 from a serine to asparagines or glycine. Both associations remained after correction for other treatments and potential confounders and the amino acid changes were replicated in independent samples. The 68 glycine variant has been described previously as correlating with drug resistance variant at position 65[21]. This change does not confer drug resistance itself but rather compensates for the reduced fitness from a change at position 65[22]. In agreement with this we observed significant interactions between the changes at position 68 and both 65 and 106 (Supplementary Table 3).

The second benefit of a GWAS approach was the ability to identify novel associations outside of candidate regions of the genome. Here we observed a novely associated SNP outside of the RT region of the Pol gene traditionally assumed to contain all genetic variants that provide resistance to NNRTIs. This association with failure on tenofovir (an NNRTI) was instead within amino acid 91 of the matrix protein of the Gag polyprotein. The effect remained after correction for effects of other drugs, population stratification and relatedness (Supplementary Table 4). This variant leads to a change in amino acid from the reference arginine to glycine, an uncommon change though the region is highly polymorphic. In comparison to the other variants (mean odds ratio of 3.70, range 1.72–11.91), the effect size was slightly smaller at 1.78 suggesting why it previously may have been unobserved.

While the current results validate the applicability of GWAS to the HIV genome, there are some limitations. As previously mentioned, not all known DR variants were identified at genome-wide significance, though given many were nominally significant, this is likely to reflect small sample size. Related to this is a bias in which types of variants were more likely to be identified in our study design. These would have related to two groups of variants. First, we would have had greater power to detect drug resistance variants that also reduce viral fitness, meaning they would only exist at high frequencies when directly under selection from treatment. Second, our study design would favour identifying variants that had effects specific to one drug rather than a class of drugs, due to most samples having been exposed to at least one drug from each class. This was a result of the now widespread usage of ART by infected individuals and subsequent focus of sequencing efforts on treatment resistance. Lastly, we note that unlike bacterial GWAS[7], we did not observe dramatic genome-wide inflation in test statistics. Our comparison of lambda values using permuted and unadjusted p-values suggested that Bonferroni adjustment for multiple corrections is likely over conservative, while permutation adjustment may not correct for all inflation. However, analysis of principle components suggested the genome-wide associations were not confounded by geographic and genetic population structure.

Overall, our results provide a clear proof of concept on the use of GWAS within HIV and other viruses whole genome sequence data. The smaller genome size, compared to humans, means that substantially smaller samples were needed to identify associated variants. Power is also greater because sequencing allows one to test the association with the causal variant, rather than the proxy SNPs often used in human GWAS to capture several nearby correlated SNPs. With a larger percentage of the genome transcribed there should also be a larger proportion of functionally relevant variants. Additionally, viruses can themselves be used as model organisms and can be genetically modified, allowing for functional validation of identified variants in a way that cannot be performed in humans. However, these benefits of performing GWAS within viruses should not ignore the valuable lessons from human genomics, especially the need to quickly establish large sample sizes through internationally collaborative research (see [25, 26]). A focus on setting up standardised quality control pipelines, making GWAS results publically available in the form of SNP summary statistics, and pooling samples into mega-analyses (rather than meta-analysing separate studies) should be the aim of those groups generating HIV and other virus genomes.

## Methods

### Sample description

The study sampled 319 HIV-infected adults and 24 children on ART with virological failure in the Hlabisa HIV Treatment and Care Programme in South Africa for which a whole genome of HIV-1 was produced. The inclusion criteria were: ART regimen for at least 12 months followed by virological failure, defined as one viral load >1000 copies/ml. Exclusion criteria were: prior use of nucleoside reverse transcriptase inhibitor (NRTI) monotherapy or dual therapy (not including regimens for the prevention of mother-to-child transmission (pMTCT)). All individuals were seen by a physician, who performed a clinical evaluation and obtained written informed consent for the study. A 5 ml EDTA whole blood sample for HIV DR genotyping was collected during the clinical evaluation. Basic clinical and demographic data, including GPS data on household location, were collected on a clinical form and clinical and treatment information was compared with the records in the Africa Centre’s ART Evaluation and Monitoring System (ARTemis), an operational database holding treatment and laboratory monitoring information from the national ART programme in South Africa. The clinical information was entered in anonymised form into a relational sequence database, the SATuRN REGA database[27]. Further details of the study have been described previously[28, 29].

### Ethics Statement

The study was approved by the Biomedical Research Ethics Committee of the University of KwaZulu-Natal (ref. BF052/10) and the Health Research Committee of the KwaZulu-Natal Department of Health (ref. HRKM 176/10). South African legal guidelines define a person able to give informed from consent from age 17. Written informed consent was obtained from all the study participants and their parent or legal guardian in the case of paediatric patients (≤ 16 years).

### Drug exposure data

The median duration of ART among patients in this cohort was 42 months (IQR 32–53). The most common first line ART regimens were: tenofovir/stavudine/zidovudine +Lamivudine +efavirenz/nevirapine. The most common second line ART regimen were: Lopinavir (+ Ritonavir), Lamivudine, zidovudine/tenofovir. The median duration of antiretroviral failure was 27 months (IQR 17–40 months). Details on drug exposure data and DR results have been described previously [28, 29]. Drug exposure was defined by exposure at any time point prior to sequencing. Table 1 provides a basic description of the characteristics of the 343 individuals with viral WGS data included in the analysis.

### RNA extraction, PCR amplification and whole genome Sequencing

RNA was extracted from samples using the manual QIAamp Viral RNA Mini Kit (Qiagen). The near complete HIV-1 genome was amplified by a previously described RT-PCR strategy with primers modified to be more subtype C specific (Danaviah et al. CROI 2015; Abstract). The amplification involved the production of four overlapping genetic fragments of lengths of 1.9kb, 3.6kb, 3.0kb and 3.5kb. This included all nine open reading frames and partial regions of the 5′- and 3′-LTR. The DNA concentration of individual amplicons was quantified using the Qubit sdDNA HS Assay Kit (Thermo Fischer Scientific-Life Technologies). Pooled amplicons were prepared for sequencing using the Nextera XT DNA Sample Preparation kit (Illumina) and the Nextera XT DNA Sample Preparation Index Kit (Illumina), following the manufacture’s protocol. The runs comprised pools of 96 samples that included three controls (one negative sample, one inter-run and one intra-run control). All processes to generate WGS were undertaken locally at the Africa Centre laboratory, Nelson R Mandela Medical School, University of KwaZulu-Natal, South Africa.

### Bioinformatics pipeline: Whole genome quality control, assembly and phylogenetic analysis

Fastq quality control was performed using FASTQC(0.11.3) and QUASR(3.1) software applications. Reads of less than 100bp in length and a quality score lower than 30 were excluded. In addition, the reads were trimmed up to 10bp from 5′ and 30bp at the 3′ to exclude poor quality sequence at the beginning and end of reads. We noticed that the second pair read of the Illumina Nextera XT was of lower quality and that excluding the last 30bp increased quality score to > 33. We imposed these exclusion criteria in order to decrease the probability of ambiguous read mapping, which occurs when shorter reads of lower accuracy are included in assemblies [30]. Following these quality control steps, we mapped reads against a subtype C reference sequence (AF411967) with five assembly iterations using Geneious 8 (http://www.geneious.com)[31]. After assembly, we exported the data as BAM files and exported contigs as FASTA files.

In order to determine if there was clustering of sequences (i.e. sequences that were very similar with low genetic diversity), we aligned all of the whole genomes with a reference dataset for HIV-1 subtype C. The tree was constructed with HKY+Gamma site rate variation in a MPI version of RaxML. Reliability of internal nodes was evaluated by 100 bootstrap replicates. Phylogenies were analysed using Phylotype software application[32] in order to detect any clustering of sequences with high bootstrap values (>90%) and low sequence diversity (<3%). This was performed to identify pairs of closely related HIV sequences that might confound the analysis and test the sensitivity of the results to their inclusion.

### Variant calling and GWAS software adaptation

The processing of WGS data to the performing of GWAS is outlined in Figure 1, with comparison to human GWAS steps. BAM files were converted to VCF format variant calls individually for each sequence in PILON [33]. A threshold of a depth of 50 reads per base was used for a variant to be called.

As GWAS software was originally designed for diploid organisms (i.e. those with two chromosomes and so two copies of any given loci), each sample can be called either as homozygous for an allele (e.g. AA or TT) or as heterozygous (e.g. AT). While heterozygosity is incorrect in the sense that HIV is haploid, it captures an important reality of viral infection: genetic differences within the host’s viral population. We wanted to retain the feature of diploidy to account for samples with diversity at a given DR loci. We expected heterozygous samples to have an intermediate effect size compared to samples where the DR variant was either entirely non-existent or fixed. The downside of this approach was that given numerous sequence reads for each loci, some variation is expected due to sequencing errors. To account for this, we allowed for diploid calling in the following manner. If the reference allele frequency was present in >85% of reads at a loci, the loci was called as homozygous for the reference allele. A heterozygous call with one copy of the reference variant and one of the non-reference variant was made if the reference allele frequency was between 85% and 15% of reads. Finally, a homozygous non-reference call was made if the reference allele frequency was found in less than 15% of reads. While these cut-offs are simply defaults of the software, this worked as a crude calling approach for whether an individual sample’s HIV population was fixed or mixed for any given loci.

VCFs were then merged in GATK[34], then the combined VCF read into PLINK1.09[35] for GWAS analysis. Prior to analysis, several QC steps were performed. First, where multiple alleles occurred at the same loci, the reference variant and the most common non-reference variant were used to make the loci bi-allelic. Second, a minor allele frequency of greater than 1% was required for all variants. Lastly, we did not implement a restriction on missingness of data. In human GWAS, high missingness for a SNP or individual may reflect poor quality genotyping. However, in HIV WGS sequencing quality is not homogenous across the genome (see Supplementary Figure 2). As we had restricted analysis to calling variants at loci with a depth of 50 or greater, higher missingness was expected. Missingness for SNPs significantly associated with DR is reported in Table 2.

### Statistical analysis

A logistic regression was performed in PLINK1.09[35] with drug exposure as the binary outcome and each SNP as a predictor with an additive effect. All samples exposed to a given drug were compared to all that were not. To determine genome-wide significance we performed 10,000,000 permutations within PLINK1.09 both on a single SNP and genome-wide level using the --mperm command. This was performed to account for correlation between nearby SNPs which would have made Bonferroni correction for the raw number of statistical tests overly conservative. Given the smaller number of variants compared to a human GWAS, permutation using 10,000,000 for the empirical p-values was computationally feasible. As the negative correlations in the prescribing of these drugs existed, associations with the same SNP were seen in multiple analyses. However it was possible to identify when exposure was associated with the non-reference sequence (i.e. odds ratio>1) and so, presumably, which association identified the true drug resistant variant. Principal components were generated in GCTA[36].

### Replication

Genome-wide significant SNPs within the Pol region were able to be taken forward for replication in a publically available independent sample. This was the Stanford University HIV Drug Resistance Database[19, 20], where information on amino acid frequencies were available for sequences exposed to different drugs. This analysis was restricted to the 13,676 subtype C sequences. Additional analyses also made use of a subset of all publically available subtype C WGS (n=505) from the Los Alamos HIV Sequence Database (http://www.hiv.lanl.gov/). This was done to ensure our variant frequencies were in agreement with those observed elsewhere.

### Data access

Summary statistics for all SNPs of each GWAS are available online (https://figshare.com/articles/PLOSONE_DR_GWAS_HIV/3569766). Access to the full genomes of HIV can be done by application of a proposal to PANGEA_HIV (http://www.pangea-hiv.org/).

### Funding Disclosure

Research supported by a South African MRC Flagship grant (MRC-RFA-UFSP-01-2013/UKZN HIVEPI) and the Bill and Melinda Gates Foundation (BMGF) PANGEA_HIV grant. Funding for the Africa Centre’s Demographic Surveillance Information System and Population-Based HIV Survey was received from the Wellcome Trust, UK (grant 082384/Z/07/Z). The funders had no role in study design, data collection and analysis, decision to publish, or preparation of the manuscript.

## Acknowledgements

We thank all the patients for their continued support, all Africa Centre staff who contribute to maintaining ACDIS and the laboratory.

